# Rotary jet-spun porous microfibers as scaffolds for stem cells delivery to central nervous system injury

**DOI:** 10.1101/239194

**Authors:** Laura N. Zamproni, Marco A.V.M. Grinet, Mayara T.V.V. Mundim, Marcella B.C. Reis, Layla T. Galindo, Fernanda R. Marciano, Anderson O. Lobo, Marimelia Porcionatto

## Abstract

Transplanting stem cells into the central nervous system is a promising therapeutic strategy. However, preclinical trials of cell-based therapies are limited by poor local cell engraftment and survival. Here, we present a polylactic acid (PLA) scaffold to support delivery of mesenchymal stem cells (MSCs) in a mouse model of stroke. We isolated bone marrow MSCs from adult C57/Bl6 mice, cultured them on PLA polymeric rough microfibrous (PLA-PRM) scaffolds obtained by rotary jet spinning, and transplanted into the brains of adult C57/Bl6 mice, carrying thermocoagulation-induced cortical stroke. Interleukins (IL4, IL6 and IL10) and tumor necrosis factor alfa (TNFα) expression levels in the brain of mice that received PRM were similar to untreated. MSCs transplantation significantly reduced the area of the lesion and PRM delivery increased MSCs retention at the injury site. We conclude that PLA-PRM scaffolds offer a promising new system to deliver stem cells to injured areas of the brain.

**GRAPHICAL ABSTRACT:** 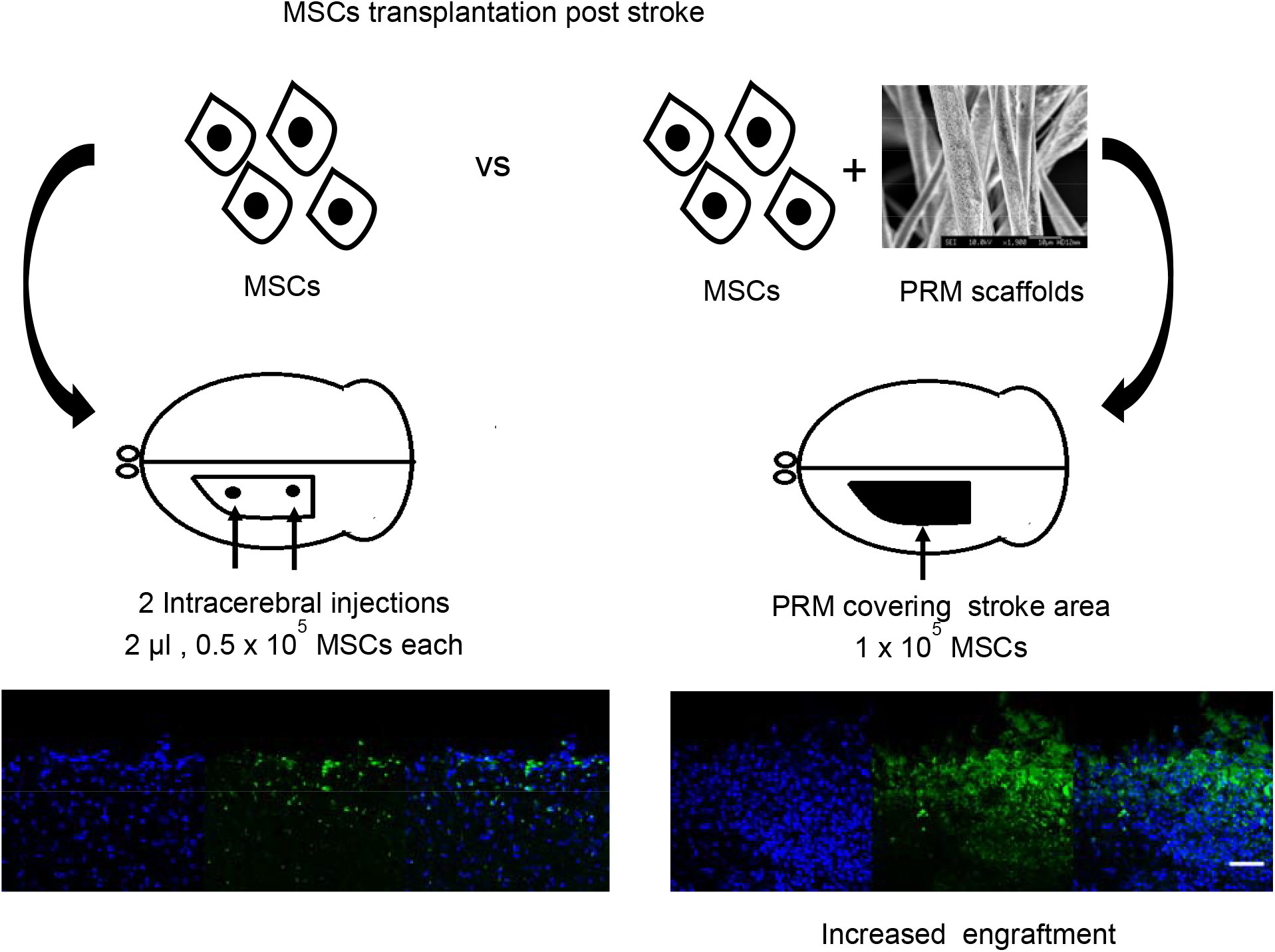

Synthetic scaffolds offer an alternative to optimize stem cell transplantation at sites of brain injury. Here, we present a rotary jet spun polylactic acid (PLA) polymer used as a scaffold to support delivery of mesenchymal stem cells (MSCs) in a mouse model of stroke. Transplantation of MSCs isolated or cultured on PRM significantly reduced the area of the lesion and PRM delivery increased MSCs retention at the injury site.

## Background

Stroke is one of the leading causes of death and severe disability in adults (1). Recombinant tissue plasminogen activator is the only FDA (Food and Drug Administration) approved treatment that can be used up to 4.5 hours after the onset of ischemic stroke. After that time frame has passed, there are no effective treatments available besides rehabilitation (2).

Stem cell-based therapies are promising for the treatment of stroke and have been extensively investigated (3–5). However, when stem cells are systemically administered, only a few cells reach the brain despite the high number of cells delivered. Delivery of MSC either via intravenous or intracardiac administration to a traumatic brain injury model in rats, showed that <0.0005% of the cells injected were found at the injury site after 3 days (6). Thus, the distribution and survival of transplanted cells are still major challenges in stem cell-based therapies and must be addressed (3).

Intracerebral (IC) administration of cells directly into the lesion cavity could be an option to overcome those issues (7). However, stroke pathophysiology involves a complex and dynamic process, including degradation of extracellular matrix (ECM) proteins, such as laminin (LMN), fibronectin, and collagens I/III and IV, offering a major obstacle to central nervous system (CNS) repair (3). It has been described a significant loss of the ECM, in addition to the loss of neurons and glia, after stroke. The damaged area is unfavorable for cell survival, resulting in a severe loss of grafted cells after transplantation (8).

Polymeric scaffolds have been used for CNS regeneration due to their ability to physically support the infiltration of host cells and to deliver exogenous stem cells locally to the site of injury (9). The use of scaffolds improves stem cell survival when the cells are delivered to an intact brain region adjacent to the lesion (7). Rotary jet spinning (RJS) offers an approach for the production of porous and bioreabsorbable polymeric microfibers for such use (10, 11). Using RJS, polymeric solutions are easily extruded during high rotation to fabricate microfibrous scaffolds, and polylactic acid (PLA) was approved by the FDA for the production of bioreabsorbable scaffolds for tissue engineering applications (12).

To date, novel cell delivery methods, such as RJS polymeric microfiber implantation, may provide the structural support required for stem cell survival, proliferation, engraftment, and differentiation after stroke (13). To the best of our knowledge, rough and microfibrous (PRM) scaffolds produced by RJS do not appear to have been evaluated as supportive structures for stem cell culture and transplantation in an *in vivo* animal model of stroke. Here, we present a method of stem cell transplant using PRM scaffolds produced by RJS for stem cell delivery directly into ischemic mouse brain cortex after injury by thermocoagulation. PRM scaffolds did not exert any deleterious effects in the mice brain cortex, and MSCs transplantation significantly reduced the area of the lesion. Additionally, PRM cell delivery increased MSCs retention to the injury site.

## Material and Methods

### 1. Microfiber production

Production of microfibers was carried out using an adapted RJS apparatus. First, 0.4 g of PLA (pellets, 2003D, Natureworks, Minnetonka, USA) were dissolved in 50 mL of dichloromethane in a closed system by magnetic stirring for 2 h. The polymeric solution was placed into a cylindrical reservoir (6 mL, 0.3 mm-diameter orifices) and RJS using a rotary tool (FERRARI MR30K) at 8,000 rpm for 30 min in a collector (placed 10 cm from the rotatory tool). We analyzed the morphology of the fibers using scanning electron microscopy (Zeiss, EV aO MA10, Oberkochen, Germany). PRM scaffolds were sterilized by immersion in 70% alcohol for at least 3 h.

### 2. Laboratory animals

The maintenance and care of the mice used in this study were performed in accordance with international standards on the use of laboratory animals. The protocols used in this study were reviewed and approved by the Committee of Ethics for the Use of Laboratory Animal in Research from UNIFESP (CEUA 8368221013). Female GFP (green fluorescent protein) positive, C57/Bl6 mice from the Laboratory for Animal Experimentation (LEA/INFAR/UNIFESP) were kept in isolated units at a room temperature of 20 ± 2°C, relative humidity of 50%, and circadian cycle of 12h intervals (light; dark). An effort was made to minimize animal suffering and reduce the number of animals used in the study.

### 3. MSCs isolation and culture

Bone marrow MSCs were isolated as previous described (14). Forty-five-day-old mice were euthanized by anesthesia with ketamine hydrochloride (90 mg/kg) (Syntec, São Paulo, Brazil) and xylazine hydrochloride (10 mg/kg) (Ceva, São Paulo, Brazil), followed by cervical dislocation; the tibias and femurs were dissected, and their epiphyses were cut. Two milliliters of Dulbecco’s Modified Eagle’s medium (DMEM, Invitrogen, San Francisco, USA) were injected into one end of the bone using a syringe (26 G; 13 mm/24, 5 mm, BD Biosciences, Franklin Lakes, USA), and the bone marrow was collected from the other end of the bone. The cell suspension was centrifuged at 400 × *g* for 10 min at room temperature. The cell pellet was suspended in 1 mL DMEM containing 10% fetal bovine serum (FBS; Cultilab, Campinas, Brazil) and 1% penicillin/1 % streptomycin (Gibco, San Francisco, USA). Cells were counted in a Neubauer chamber, resuspended in DMEM containing 10% FBS at a density of 5 x 10^6^ cells/mL, and incubated at 37 °C in a 5% CO2 atmosphere. The exchange of culture media was done every 72 h to remove nonadherent hematopoietic cells. When cells reached 90% confluence, they were trypsinized (Cultilab) in 10 mM PBS for subculture. After the second subculture, adherent cells were considered MSCs. For transplantation, 1 x 10^5^ MSCs from the 6^th^ to 10^th^ passages were seeded onto PRM 24 h prior to surgery.

### 5. Analysis of cell viability

MTT [3-(4,5-dimethylthiazol-yl)2,5-diphenyltetrazolium bromide, Sigma, St Louis, USA] assay was performed to evaluate MSCs viability after a 7-day culture over the scaffolds. 1x10^4^ cells were cultured in the bottom of a 96-well plate or over scaffolds. Culture medium (270 μL) and the MTT solution (30 μL; 5 mg/L) were added to each well of a 96-well plate, and the cells were incubated for 3 h at 37 °C. The MTT solution was aspirated, and formazan was dissolved in 180 μL dimethyl sulfoxide (DMSO, MP Biomedicals, Solon, USA). The plate was shaken for 15 min and optical density was read at 540 nm on an ELISA plate reader (Labsystems Multiskan MS, Helsinki, Finland).

### 6. Immunocytochemistry

MSCs were cultured on glass coverslips or PRM scaffolds for 48 h in DMEM containing 10 mM BrdU (5-bromo-2’-deoxyuridine, Sigma, St. Louis, USA). The cells were then fixed by 4% paraformaldehyde (PFA), permeabilized with 0.1% Triton X-100 (Sigma), and immunostained with Alexa Fluor^®^ 594-conjugated anti-BrdU and FITC-conjugated phalloidin (Molecular Probes, Eugene, USA). Cells were analyzed by scanning confocal microscopy (TCS, SP8 Confocal Microscope, Leica, Wetzlar, Germany).

### 7. Evaluation of apoptosis by TUNEL (terminal deoxynucleotidyl transferase-mediated dUTP nick end labeling) assay

MSCs were cultured on glass coverslips or PRM scaffolds, and analyzed after 7 days *in vitro*. TUNEL assay was conducted following the protocol suggested by the manufacturer (Kit S7111, Millipore, Darmstadt, Germany). Cells were analyzed by scanning confocal microscopy (TCS, SP8 Confocal Microscope).

### 8. ELISA

1x10^5^ cells were cultured in the bottom of a 24-well plate or over scaffolds. After 24 hours the media was changed. Conditioned media was collected 48 hours later. CXCL 12 in conditioned media was quantified using Mouse CXCL12 DuoSet ELISA (R&D systems, Minneapolis, USA).

### 9. Mouse model of ischemic stroke

Stroke was induced by thermocoagulation in the submeningeal blood vessels of the motor and sensorimotor cortices. The thermocoagulation protocol was adapted for mice from previously described protocols for rats (15). Briefly, animals were anesthetized with ketamine hydrochloride (90 mg/kg) and xylazine hydrochloride (10 mg/kg), and placed in a stereotaxic apparatus. Stereotaxic coordinates were established from bregma (anterior +2, lateral +1, posterior -3) to localize the frontoparietal cortex (Supplementary Figure 1). The skull was exposed, and a craniotomy was performed, exposing the left frontoparietal cortex. Blood was thermocoagulated transdurally by the placement of a hot probe in the dura mater. The skin was sutured, and the animals were kept warm and returned to animal facility after recovery from anesthesia.

Animals were randomly separated and assigned to the following experimental groups: Control: no intervention; Lesion: thermocoagulation; Lesion+PRM: thermocoagulation and the addition of PRM scaffolds; Lesion+PRM+MSC: thermocoagulation and the addition of PRM seeded with MSCs; Lesion+MSC: thermocoagulation and intracerebral injection of MSCs (1 x 10^5^ cells in DMEM 4 μl, injected in 2 points: anterior +1, lateral +1.5, ventral -1 and posterior -2, lateral + 1.5, ventral -1). All cells used for transplant procedures were obtained from GFP expressing animals.

### 10. Euthanasia and histological analysis

Mice were euthanized by lethal anesthesia 12 or 30 days after injury and intracardially perfused with 100 mM PBS followed by 4% PFA in 100 mM PBS (pH 7.4). Next, the brains were removed, immersed in 4% PFA for 24 h, and cryopreserved in PBS containing 30% sucrose for 48 h. To estimate the amount of cerebral tissue lost after stroke, the brains were sectioned into 1 mm coronal slices using a manual sectioning block, and the lost tissue area in each slice was determined by subtracting the area of the injured hemisphere from the area of the normal hemisphere using Image J software. The volume of brain tissue lost was determined the sum of each slice area multiplied by the thickness (1 mm): lost area = Σ (area of contralateral side – area of ipsilateral side) x 1 (16). For histological analyses, brains were sectioned into 30 μm coronal slices at −20 °C using a CM 1850 cryostat (Leica).

### 11. Immunohistochemistry

For immunofluorescence staining, cortical sections were incubated overnight at 4°C with anti-GFP (1:100, rabbit IgG, Merck Millipore), anti-glial fibrillary acidic protein (GFAP, 1:1000, chicken IgG, Merck Millipore) or anti-IBA1 (1:100, rabbit IgG, Merck Milipore). After washing with PBS, the sections were incubated at room temperature with the appropriate secondary antibodies conjugated to Alexa Fluor^®^ 488 or 594 (1:500, Invitrogen). Nuclei were stained with DAPI (1:500, Molecular Probes). Glass slides were mounted using Fluoromount G (Electron Microscopy Sciences, Pennsylvania, USA). The fluorescently labeled tissue slices were analyzed using a scanning confocal inverted microscope (TCS, SP8 Confocal Microscope), and image overlays were generated using ImageJ software, version 5.01. Stained cells were quantified by number of cells by mm^2^. Data from 3 animals in each group, 3 sections per animal were analyzed.

### 12. Quantitative PCR (qPCR)

Total RNA was isolated from ipsilateral stroke cortex using the Trizol^®^ reagent (Life Technologies, Thermo Fisher Scientific, Wilmington, MA, USA), and RNA concentrations were determined using a NanoDrop ND-1000 instrument (Thermo Fisher). Reverse-transcriptase reactions were performed with the ImProm-II Reverse Transcription System (Promega, Madison, WI, USA) using 2 μg total RNA. qPCR was performed using Brilliant^®^ II SYBR^®^ Green QPCR Master Mix (Applied Biosystems, Thermo Fisher Scientific) and the Mx3000P QPCR System; MxPro qPCR software was used for the analysis (Stratagene, San Diego, CA USA). Primers sequences are shown in Table 1.

**Table1:**
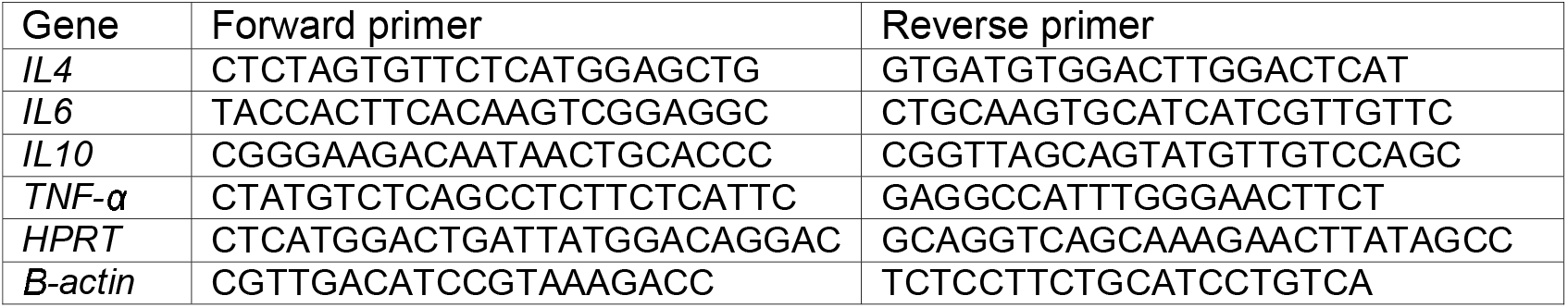
Primers sequences.

Values are expressed relative to those from the control group. For quantification, the target genes were normalized using an endogenous control gene *Hprt* or *Bactin*. The threshold cycle (Ct) for the target gene and the Ct for the internal control were determined for each sample, and triplicate experiments were performed. The relative expression of mRNA was calculated using the 2^−ΔΔCt^ method (17).

### 13. Statistical analysis

The statistical significance of the results was determined using the Student’s *t* test or oneway ANOVA plus Tukey’s method for multiple comparisons. The results are expressed as the average ± standard error. A *p* value ≤0.05 was considered statistically significant.

## Results

### Characterization of rough microfibers: Alignment, size, and cytocompatibility

We synthesized PRM scaffolds using PLA. Aligned PRM obtained had a 3.93 ± 2.30 μm diameter and pores of 0.57 ± 0.15 μm (Figure 1A).

**Figure 1:**
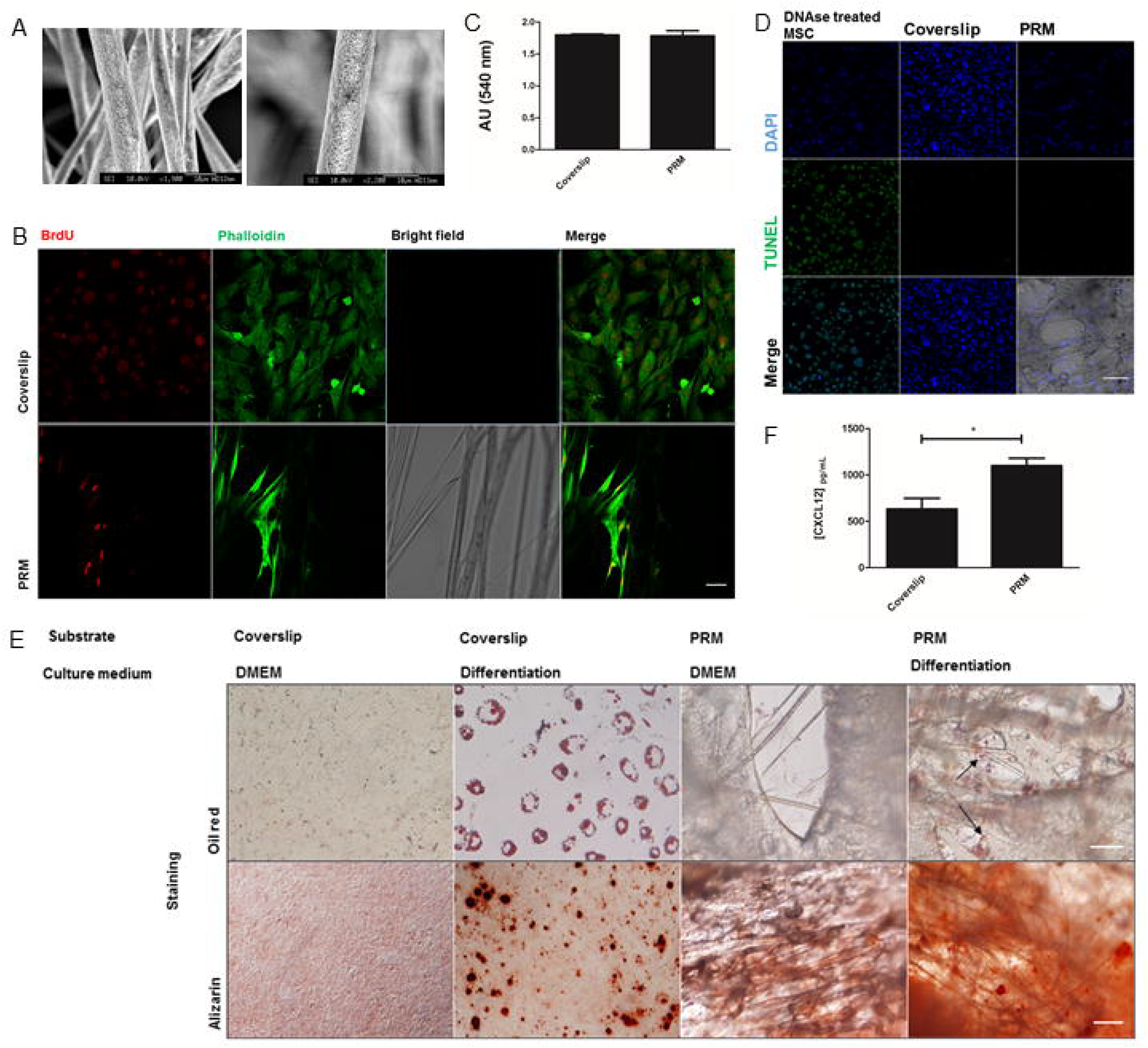
MSC compatibility with PRM scaffolds. (A) Characterization of PRM scaffolds by scanning electron microscopy. (B) BrdU and phalloidin immunostaining of MSCs on coverslips or PRM scaffolds. Cells were able to adhere and incorporate BrdU into DNA (proliferating cells) in either condition (n = 2, experiment in triplicates). Scale bar = 100 μm. (C) After seven days in culture, MTT assay showed similar survival of MSCs when cells were cultured on coverslips or PRM scaffolds (p > 0.05, Student’s *t* test, n = 5). (D) TUNEL assay showed that MSCs cultured on coverslips or PRM scaffolds did not undergo apoptosis (n = 2, experiment in triplicates). Scale bar = 100 μm. (E) MSCs cultured on coverslips or PRM scaffolds were induced to differentiate in osteocytes and adipocytes. After 3 weeks, cells were stained with Oil Red (staining of lipid vesicles in red) or Alizarin Red (staining of calcium deposits in red). MSCs cultured on PRM scaffolds preserved multipotency (n = 2, experiment in triplicates). Upper scale bar = 50 μm; lower scale bar = 200 μm. (F) PRM induced a 50% increase in CXCL12 secretion by MSCs (*p* < 0.05, Student’s *t* test, n = 5). MSCs: mesenchymal stem cells, PRM: polymeric rough microfibers, BrdU: 5-bromo-2’-deoxyuridine, MTT: 3-(4,5-dimethylthiazol-yl)2,5-diphenyltetrazolium bromide, TUNEL: terminal deoxynucleotidyl transferase-mediated dUTP nick end labeling, DAPI: 4’,6-diamidino-2-phenylindole.

We evaluated cytocompatibility of PRM scaffolds with MSCs using five different approaches. First, we observed integrity of cell morphology by phalloidin-stained the cytoskeleton (Figure 1B). Proliferation, survival and cell death were assessed by BrdU incorporation (Figure 1B), MTT (Figure 1C), and TUNEL (Figure 1D), respectively. Clearly, MSCs proliferated when cultured on the PRM scaffolds, and were able to spread, similar to those observed when MSCs are grown on glass coverslips (Figure 1B). After seven days, the number of viable cells was similar between cells plated on either the PRM scaffolds or coverslips (Figure 1C). We did not identify apoptotic nuclei in the growing MSCs (Figure 1D), confirming the MTT results that MSCs remain viable when cultured on PRM scaffolds for up to seven days. Finally, we analyzed the multipotency of MSCs when cultured on the scaffolds by inducing adipogenic and osteogenic differentiation by staining the cells with either Oil Red (stains lipid vesicles in adipocytes) or Alizarin Red (stains calcium deposits produced by osteocytes). The results show that PRM scaffolds did not alter MSCs multipotency (Figure 1E).

Since CXCL12 is key participant in MSCs retention to sites of injury, we investigated CXCL12 levels in the medium when MSCs were cultured on PRM and compared to CXCL12 levels when MSCs were cultured in 24-well plate. We found that PRM induced a 50% increase in CXCL12 secretion by MSCs (Figure 1F).

### PRM are safe for brain implantation

Despite good *in vitro* results, our main concern was that PRM could be perceived as a foreign body, and could enhance damage to the brain due to increased immune response. In order to investigate if PRM elicited any adverse immune response, we implanted PRM in the brain of mice that underwent thermocoagulation-induced stroke, and were euthanized 12 or 30 days later. In all brains, we found PRM scaffolds firmly attached to the skull at the site of injury, and was removed together with the skull (Figure 2A). The brain showed a necrotic patch that was easily detached from the normal parenchyma, leaving an atrophic area (Figure 2B). Coronal sections of 1 mm thick were obtained, and the volume of brain parenchyma lost 12 and 30 days after injury was measured. In both endpoints, brain of mice that received the scaffolds lost similar volume of parenchyma as those that did not receive the scaffolds (Figure 2C).

**Figure 2:**
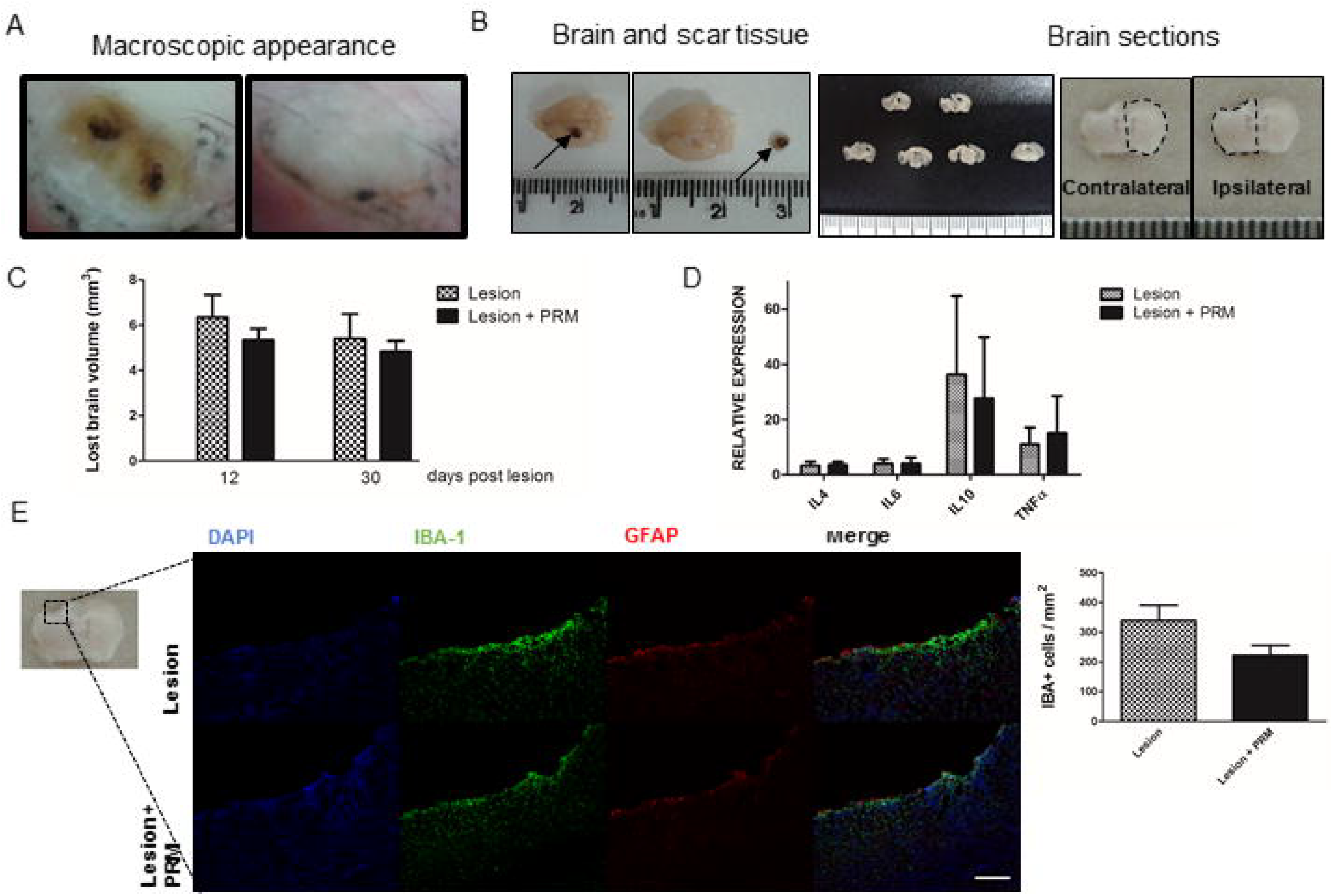
*In vivo* evaluation of PRM safety. (A) Macroscopic findings at euthanasia 30 days post induction of stroke. In mice who received PRM, they were found at the craniectomy area, firmly attached to the skull bone (right panel). (B) Brain lesion assessment. The necrotic area easily detaches from the normal brain parenchyma (arrows). The brain was sliced in 1 mm sections and the volume of the necrotic area of each slice was obtained by subtracting the area of the injured hemisphere (ipsilateral) from the area of the normal hemisphere (contralateral), multiplied by the thickness of the slices. (C) Comparison of lost brain volume 12 and 30 days after thermocoagulation-induced stroke. PRM implanted at the stroke site did not increase necrotic area (n = 3-5, *p* > 0.05, Student’s *t* test). (D) Relative expression levels of interleukins and TNFα the injury site 12 days after stroke induction showed no statistically significant difference (n = 3, *p* > 0.05, Student’s *t* test). (E) The number of microglial cells (IBA1+) surrounding the injured area showed no statistically significant differences. Three representative large field photos from each animal were analysed (n = 3, *p* > 0.05 Student’s *t* test). Scale bar = 200 μm. MSCs: mesenchymal stem cells, PRM: polymeric rough microfibers, IL: interleukin.

Previous studies showed that CNS implants induce an increase in IBA1+ cell population, peaking between 1 and 2 weeks after implantation, and slowly disappearing later (18). For that reason, we quantified the expression of inflammatory cytokines and IBA-1+ cells that infiltrated 12 days post stroke induction. PRM scaffolds did not interfere with IL4, IL6, IL10, and TNFα expression levels in the brain cortex adjacent to the lesion, neither increased microglial infiltrates into the lesion (Figures 2 D and E). These results suggest that PLA-PRM scaffolds do not increase immune response.

### MSC transplantation decreases the lesion size and PRM increases MSC retention

After verifying PRM safety, we investigated whether the scaffolds were suitable for MSCs delivery into a brain lesion. We transplanted 1 x 10^5^ MSCs 24 h previously seeded on PRM scaffolds, in the brains of mice subjected to thermocoagulation-induced stroke and compared to mice receiving two intracerebral injections of 0.5 x 10^5^ MSCs each (Supplementary Figure 1) and analysed the outcome after 12 days.

The volume of brain parenchyma lost after stroke was significantly reduced in animals that received MSCs either directly into the injury or cultured on PRM scaffolds (p < 0.05) (Figure 3A). Interestingly, we found three times more GFP+ MSCs in the perilesional area when cells were delivered by PRM compared to administration of isolated MSCs (Figure 3B and C).

**Figure 3:**
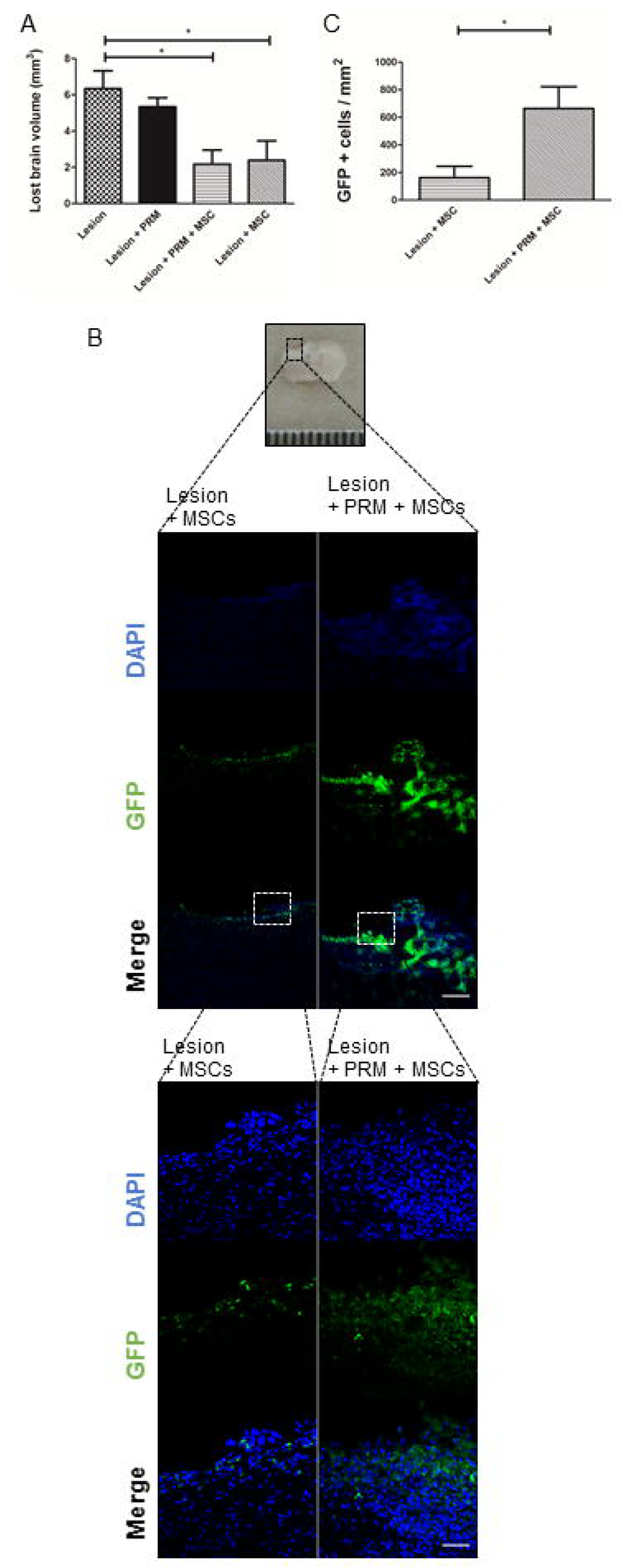
PRM increase MSCs engraftment at lesion site. (A) MSCs delivered isolated or adhered at PRM decrease the volume of brain parenchyma lost after stroke (n = 3-5, \**p* < 0.05, one-way ANOVA and Tukey’s post-test). (B) Immunolocalization of transplanted GFP+ MSCs at the lesion site. Upper panel scale bar = 200 μm. Lower panel scale bar = 50 μm. (C) The number of transplanted cells GFP+ MSCs surrounding the injured area was double when cells were transplanted adhered to PRM. Three representative large field photos from each animal were analysed (n = 3, *p* < 0.05 Student’s *t* test). MSCs: mesenchymal stem cells, PRM: polymeric rough microfibers, DAPI: 4’,6-diamidino-2-phenylindole, GFP: green fluorescent protein.

## Discussion

Transplantation of stem cells into the lesioned brain is now considered a potential treatment for brain injury and stroke. However, after stroke, a focal cavity is formed at the injury site, which fails to provide structural support for the attachment of transplanted cells. Scaffolds made of biologically compatible materials can provide better microenvironment for the survival and engraftment of transplanted cells. PLA, a known biodegradable and biocompatible polymer already approved by the FDA for medical applications (19), has been extensively studied and used in implantable devices (20, 21). Within 6 to 12 months, implanted PLA particles are absorbed and metabolized in the target tissue into lactic acid (22). The partially degraded portions of the PLA scaffolds that remain are cleared through the blood, liver, and kidneys (22).

Scaffolds should not present any biological toxicity, and induce minimal or zero immune response (23). The PRM that we present here fulfill those requisites. Besides, a microfibrous design may be useful to obtain optimal functional recovery (24). RJS has several advantages compared with other nano/microfiber fabrication methods (25). RJS is readily applicable to polymer emulsions and suspensions, allowing the use of polymers that cannot be manipulated using other techniques. High porosity, provided by the method, promotes cell adhesion, since the pores mimic the complex architecture of the ECM (26). CXCL12 and its receptor CXCR4 are key participants in MSCs homing and retention to sites of injury (27). Previous studies show that cells injected at the lesion site tend to migrate to the ventricle walls probably attracted by the high CXCL12 levels in the neurogenic niche at the sub ventricular zone (28, 29). We found that MSCs in PRM secrete up to 50% more CXCL12 than in coverslips. That finding could represent a possible advantage in using PRM for transplanting MSCs, since higher local levels of CXCL12 could facilitate cell retention at injury site and offers a possible explanation to the greater number of transplanted MSCs found at lesion site when they were delivered with PRM.

The major disadvantage of solid scaffolds such as the PRM that we developed here is that they must be surgically implanted which could increase morbidity due to surgical procedure risk. Despite this, patients who suffered large cortical stroke, and are not eligible for thrombolytic therapy, are often submitted to decompressive craniectomy and duroplasty with dural graft implant. Microfibers have already been tested as dural substitutes with good results (30, 31). Our idea is that this group of patients could benefit from receiving a functionalized dural graft containing stem cells. This alternative cell delivery method could be used with minimal changes in surgical procedure and improve stem cell survival due to the delivery of stem cells to an intact brain region adjacent to the lesion (7).

In this study, we propose a novel approach for stem cell delivery into brain injury. PLA-PRM scaffolds were developed and demonstrated to be suitable for the transplantation and to deliver MSCs into brain injured areas. We found no evidence that the scaffolds increased inflammation or caused no further damage to the brain. MSCs delivered via PRM scaffolds penetrated the injury site and promoted reduction of the injured area. Therefore, we propose a new method for stem cell transplantation into the brain, which is capable of improving cell engraftment at the site of injury.

## Acknowledgments

We kindly thank FAPESP (LNZ: 2013/165338; MP: 2009/05700-5, 2012/00652-5; AOL: 2011/17877-7, 2015/09697-0; FRM: 2011/20345-7, 2016/00575-1) and CNPq (MP: 404646/2012-3, 465656/2014-5) for financial support.

---

Supplementary Figure 1: Schematic drawing showing craniectomy coordinates and MSCs treatments.

MSCs: mesenchymal stem cells, PRM: polymeric rough microfibers.

